# New features of micro-RNA regulation of mRNA translation and stability revealed by expression of targeted or not targeted reporter genes

**DOI:** 10.1101/2022.03.16.484663

**Authors:** Dorota Hudy, Joanna Rzeszowska-Wolny

## Abstract

The existence of translation regulation by RNA-induced silencing complexes (RISCs) composed from Argonaute proteins and micro-RNAs is well established, however the mechanisms underlying specific cellular and miRNA effects and the way in which specific complexes arise are not completely clear.

Here we describe experiments with *Renilla* and *Firefly* luciferase reporter genes transfected on a PsiCheck2 plasmid into human cancer HCT116 or Me45 cells where only the *Renilla* gene contained or not sequences targeted by micro RNAs (miRNAs) in the 3’UTR. The effects of targeting were miRNA-specific; miRNA-21 caused strong inhibition of translation whereas miRNA-24 or Let-7 caused no change or an increase in global reporter *Renilla* luciferase synthesis, and the mRNA-protein complexes formed by reporter transcripts in both cell types differed as shown by sucrose gradient sedimentation. In both cell types the presence of miRNA targets on *Renilla* transcripts affected expression of the co-transfected non-targeted *Firefly* luciferase, and *Renilla* and *Firefly* transcripts were found in the same sucrose gradient fractions. We also observed that specific anti-miRNA oligoribonucleotides influenced expression of the *Firefly* as well as of the *Renilla* gene, suggesting modulation of non-targeted transcript expression by miRNAs. Our results indicate the existence of interactions between miRNA-regulated and -unregulated transcripts and suggest that the use of the latter as a normalizers in experiments may be biased. We also discuss some hypothetical mechanisms which could explain the observed miRNA-induced effects.

## INTRODUCTION

Regulation of gene expression includes many mechanisms modulating transcription and post-transcriptional processes, and is important to achieve and maintain the correct levels of proteins in cells. Much of the regulation takes place on messenger RNA (mRNA) during translation and post-transcriptionally, and the initiation step is critical for the regulation of translation (Hinnebusch 2011; Jackson et al. 2010; Pelletier and Sonenberg 2019; Shirokikh and Preiss 2018). A specific role in modulating the efficiency of translation of particular mRNAs is played by micro-RNAs (miRNAs), small 19-22 nucleotides-long RNAs that regulate the expression of most human mRNAs (Friedman et al. 2009). One miRNA can influence the translation of many mRNAs, and one mRNA can be affected by many miRNAs. MiRNAs interact with Argonaute proteins, of which there are four types in human cells (AGO1-4), to form the core of RNA-induced silencing complexes (RISCs) which recognize specific target mRNAs, normally in their 3’UTR, by complementary base pairing. This causes downregulation of mRNA expression by inhibition of translation or accelerated degradation of the mRNA. In human and animal cells most miRNAs require only a “seed” region of perfect complementarity with the mRNA for translation inhibition (Bartel 2009). In plant cells miRNAs which are perfectly complementary to their mRNA target cause mRNA cleavage which is rare in animal cells and in human cells only AGO2 has an RNase domain (Höck and Meister 2008).

The mechanism(s) by which miRNAs and RISC modulate translation is still not completely clear. RISC-induced degradation of mRNA depends on the type of Argonaute and the complementarity of miRNA with an mRNA target. The position of mismatches within or close to the seed sequences and cleavage site are determinant for the results of RISC action (Salomon et al. 2015).

Depending on cell type, conditions, and miRNA-mRNA interactions there are different pathways which lead to inhibition of translation initiation and/or formation by mRNA and proteins of complexes visible as cytoplasmic foci such as P-bodies or stress granules (SGs). P-bodies are large complexes containing mRNAs and proteins but devoid of translation machineries and enriched with translational repressors, factors involved in nonsense-mediated RNA decay, AU-rich element decay, miRNA silencing, and the de-capping protein Dcp1a. Stress granules have a different protein composition, containing the protein TIA1 but not Dcp1a (Leung et al. 2006). It is not clear how miRNAs influence the creation of P-bodies or stress granules. All human AGO family members are found in P-bodies and SGs; the localization of AGO2 to P-bodies is not dependent on miRNA because it still appears in these structures when miRNA biogenesis is blocked. In contrast, AGO2 localization to SG requires miRNAs (Leung et al. 2006).

Although the most common outcome of the action of miRNAs and RISC is inhibition of gene expression, interactions between miRNAs and mRNA which upregulate translation have been reported in the cases of some mRNAs for ribosomal proteins (Ørom et al. 2008) and for activation of hepatitis C virus RNA (Fehr et al. 2012; Roberts et al. 2011). Both effects were reported for transcripts with shortened poly(A), and depended on the growth conditions; interactions involving miRNA, AGO2, and FXR1 stimulated translation in quiescent cells but repressed translation in cycling cells (Bukhari et al. 2016; Truesdell et al. 2012; Vasudevan et al. 2008).

In the present work we studied another facet of regulation by miRNAs, the influence of miRNA-targeted sequences in a reporter gene mRNA on expression of other genes. We compared the regulation of expression of the same reporter genes in two different cell types and the expression of co-transfected miRNA-targeted and non-targeted reporter genes. The results suggest that the presence of miRNA-targeted sequences in one gene can influence the expression of another non-targeted gene, and that different cell types respond to the same regulatory sequence in mRNA by creation of different types of non-ribosomal complexes. We propose a model in which intermolecular interactions, including RNA-RNA interactions, may explain these effects.

## RESULTS

### miR-21, miR-24 and Let-7 influence the expression of non-targeted Firefly luciferase as well as targeted *Renilla* reporter genes

Using a common assay, we co-transfected HCT116 and Me45 cells with psiCHECK2 plasmids containing two luciferase genes, a reference *Firefly* gene and a reporter *Renilla* gene with or without eight tandem repeats of target sequences for miRNAs Let-7, miR-21 and miR-24 in the 3’UTRs. The presence of miR-21 targeted sequences in the *Renilla* luciferase gene caused some decrease in the level of luciferase mRNA, less significant in Me45 cells, and a nearly complete inhibition of translation in both cell lines. The presence of an miR-24 target was accompanied by a decrease in mRNA level but a slight increase of protein level in Me45 cells, whereas in HCT116 cells both the reporter mRNA and protein levels decreased. A Let-7 target on the transcript resulted in a decreased level of mRNA and decreased translation in Me45 cells, and did not significantly influence mRNA and increased protein levels in HCT116 cells [Fig.1].

**Figure 1.**
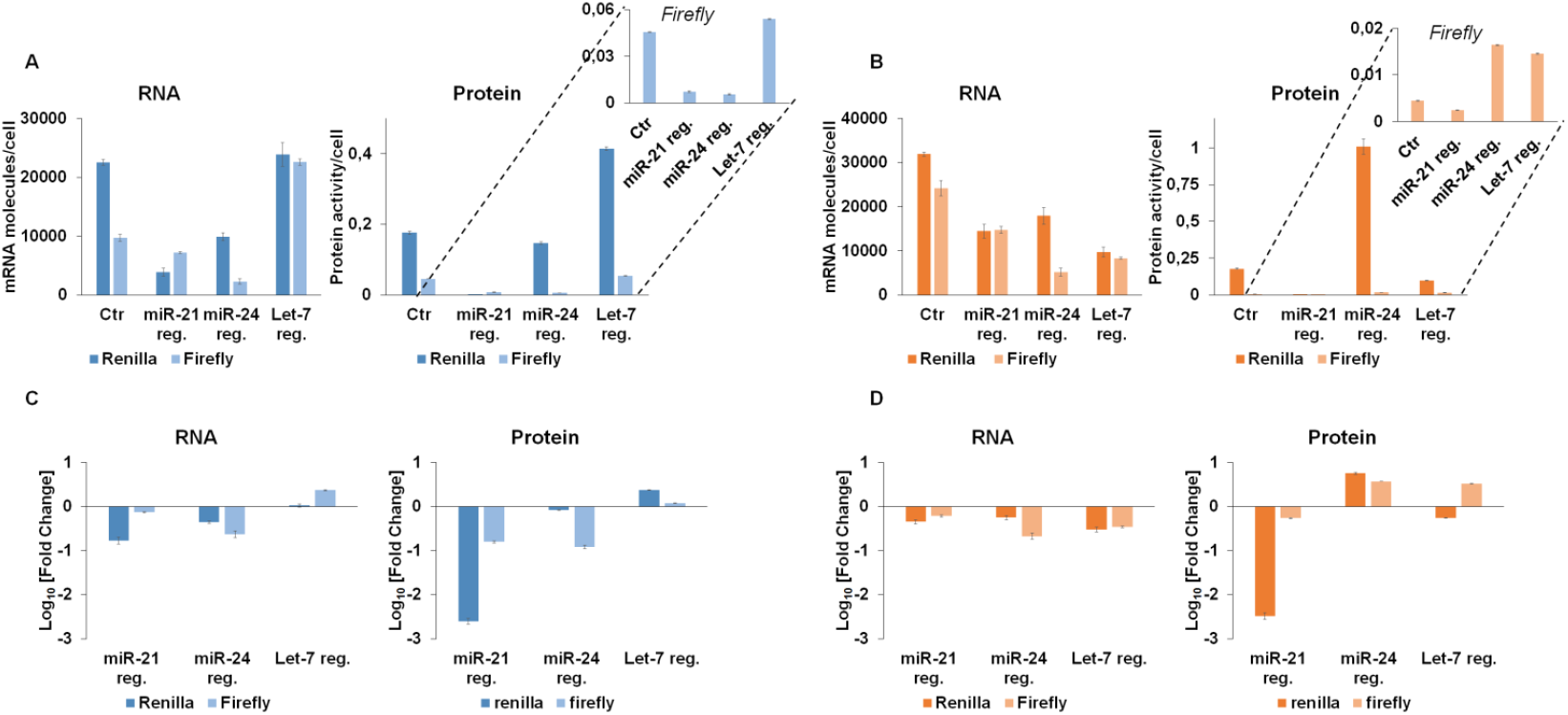
Expression of *Renilla* and *Firefly* luciferase reporter genes transfected into HCT116 (A and C) or Me45 (B and D) cells. (A) and (B) show the levels of control and miRNA-targeted *Renilla* and of non-targeted co-transfected *Firefly* luciferase mRNA transcripts (molecules/cell) and proteins (activity/cell) in HCT116 and Me45 cells, respectively. (C) and (D) show fold changes of the levels of luciferase transcripts and proteins resulting from the presence of miRNA targets in the *Renilla* gene. The results are means from three independent experiments performed on the same cell populations and vertical bars show standard deviation.

Conventionally, in similar transient transfection experiments changes in expression of the *Firefly* gene are used as indicators of transfection efficiency and the levels of its mRNA and protein serve to normalize those of *Renilla* mRNA and protein. However, in our experiments the levels of luciferase mRNA and protein (activity/cell) from the *Firefly* gene, which was not targeted by any miRNA, also changed when cells were co-transfected with a *Renilla* luciferase gene regulated by a miRNA. In some cases, for example in both HCT116 and Me45 cells, the changes of *Firefly* mRNA were larger than those for transcripts targeted by miRNA-24 (Fig1C, D, bars with lighter color). *Firefly* gene expression showed unexpected effects depending on the regulation of neighboring transcripts by miRNAs; for example, in Me45 cells regulation of the *Renilla* gene by miR-24 or Let-7 was accompanied by a decrease of *Firefly* luciferase mRNA together with an increase of its protein level (Fig. 1C and D). The decrease in mRNA could be interpreted as the result of decreased transfection efficiency, but the simultaneous increase of protein level in the same sample cannot be explained in this way and must result from a change in regulation of translation of *Firefly* mRNA.

In Table 1 we summarize the efficiencies of translation calculated as quantity of protein (activity) per molecule of mRNA for the *Renilla* luciferase gene targeted by different miRNAs and for the co-transfected, non-targeted *Firefly* luciferase genes.

**Table 1.** Translation efficiency of miRNA-targeted *Renilla* or non-targeted co-transfected *Firefly* transcripts [protein activity, a.u./mRNA molecule]*

|  |  | No target | miR-21 | miR-24 | Let-7 |
| --- | --- | --- | --- | --- | --- |
| Me45 cells | *Renilla* | 5.56±0.10 | 0.04±0.01 | 56.82±5.62 | 10.15±1.15 |
|  | *Firefly* | 0.18±0.01 | 0.16±0.01 | 3.23±0.46 | 1.75±0.06 |
| HCT116 cells | *Renilla* | 7.82±0.21 | 0.12±0.02 | 14.88±0.94 | 17.45±1.40 |
|  | *Firefly* | 4.71±0.25 | 1.01±0.05 | 2.46±0.42 | 2.39±0.05 |

\**Renilla* and *Firefly* luciferases are different enzymes which use different substrates. The arbitrary units based on luminescence calculated for each type are different and results for *Renilla* and *Firefly* cannot be compared directly.

The translation efficiency of the Renilla transcript, which does not contain any regulatory sequences in its 3’ UTR, is similar in both cell types, however one has to remember that Renilla and Firefly luciferases are different enzymes and that protein units based on luciferase activity are not compatible. Introduction of an miRNA-targeted sequence to the transcript changes the translation efficiency of both Renilla and Firefly protein. In HCT116 cells regulation of the Renilla transcripts by miRNA had less effect on the Firefly luciferase translation efficiency, but this influence was more significant in Me45 cells. The presence of miR-21-targeted sequences on Renilla transcripts is correlated with decreased translation efficiency of both Renilla and Firefly transcripts in both cell types, but targeting of Renilla transcripts by miR-24 or Let-7 was correlated with a decrease of translation efficiency in HCT116 cells but an increase in Me45 cells, suggesting that the same reporter genes are regulated by different mechanisms in HCT116 and Me45 cells.

Changes in translation efficiency calculated as the amount of protein activity per mRNA molecule could result from an increase of the amount of protein translated from the same amount of mRNA (higher speed of translation, easier translation initiation with the same number of mRNA molecules), or alternatively from a decrease in the number of mRNA molecules with the same speed of translation and level of protein (higher degradation of non-translated mRNA).

### Sucrose gradient centrifugation of complexes formed by *Firefly* and *Renilla* mRNAs

It was interesting to examine if the different effects of sequences targeted by Let-7, miR-21 or miR-24 on luciferase mRNA and protein levels in the two cell lines were related to differences in the type of complexes containing mRNA during or after its translation. To separate mRNA-protein complexes of different sizes (ribosomes, polysomes, and non-complexed mRNA) cytoplasmic extracts were centrifuged in 15% to 45% sucrose gradients which were fractionated into 100 fractions and the OD at 260 nm was measured in each fraction (Fig. 2A). These small fractions were pooled into 5 larger fractions which contained enough material to perform assays of mRNA content; fraction 1 (top of the gradient) should contain RNA free or bound to proteins, fraction 2 small ribosomal subunits and complexes of similar size, fraction 3 large ribosomal subunits and monosomes, fraction 4 light polysomes and complexes of similar size, and fraction 5 heavy polysomes and the heaviest non-ribosomal complexes (Fig.2A). The level of mRNA for *Renilla* and *Firefly* luciferases in each fraction were assayed by RT-qPCR (Fig.2B and C).

**Figure 2.**
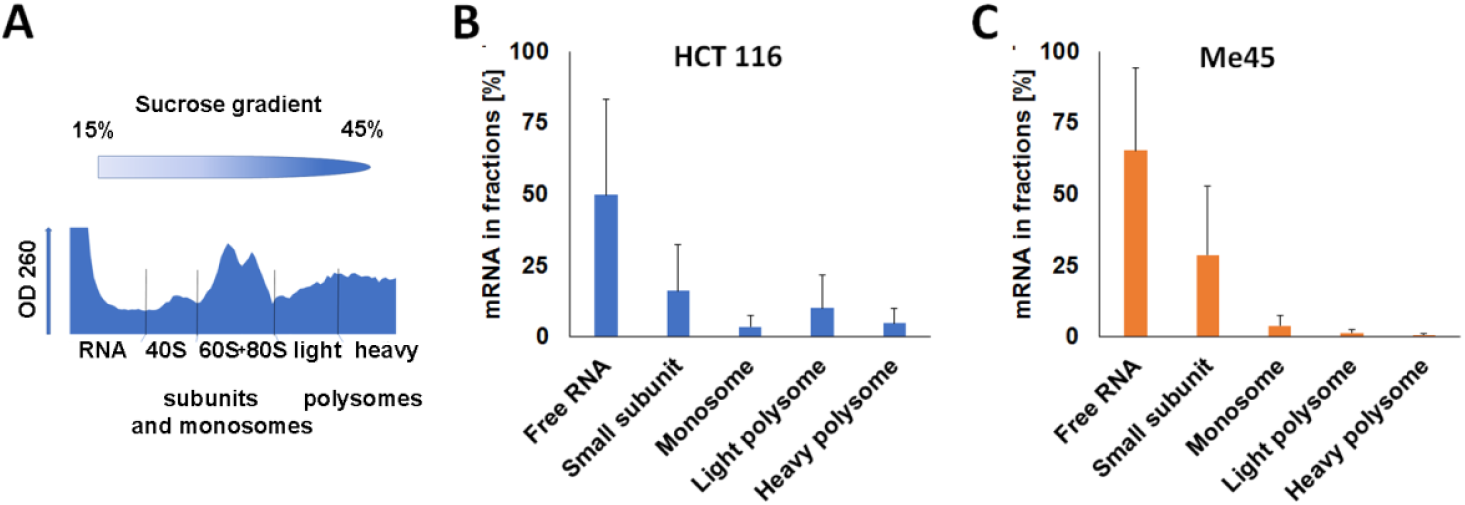
Distribution of mRNA for *Renilla* luciferase in fractions from sucrose gradients of cell extracts. (A) optical density profile of a gradient (HCT116 cells); (B, C) distribution of mRNA in sucrose gradients of extracts from (B) HCT116 or (C) Me45 cells (mean from three independent experiments, vertical lines show standard deviation).

Most of the *Renilla* luciferase mRNA was found in fraction 1 in both cell lines. The levels of reporter mRNA in the next fractions 2, 3, 4 and 5 showed a gradual decrease, except in fraction 4 from HCT116 cells which showed some increase in comparison to the preceding and following fractions. The high level of mRNA in the “free RNA” fraction was specific only for reporter gene mRNAs; transcripts for the non-reporter genes *GAPDH* and *RPL41* were distributed differently (Fig. 3).

**Figure 3.**
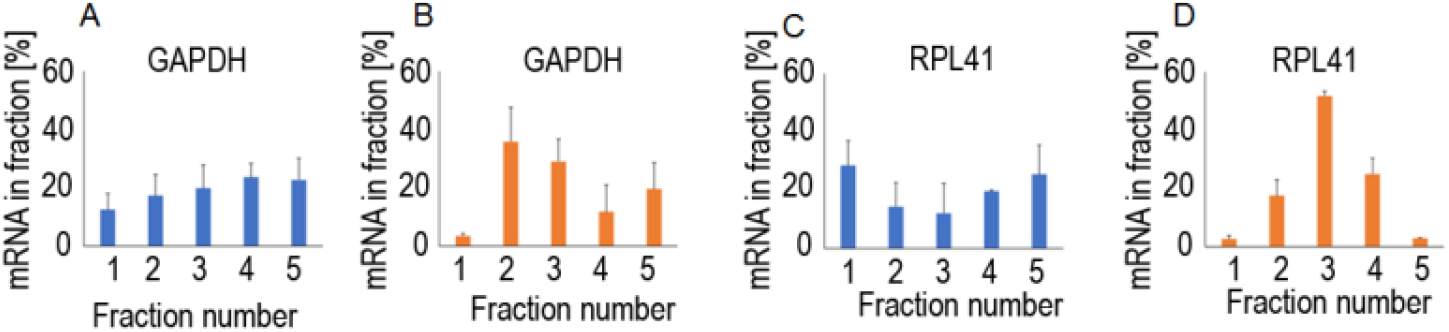
Distribution of mRNA for the *GAPDH* (A,B) and *RPL41* (C,D) genes in sucrose gradient fractions from HCT116 (A, C) and Me45 (B, D) cells.

Next, we examined the distributions of miR-21, miR-24 and Let-7-targeted *Renilla* luciferase and untargeted *Firefly* luciferase mRNAs in similar sucrose gradient fractions of cell extracts (Fig. 4). The distributions of *Renilla* luciferase mRNA were different in each cell type and depended on the miRNA-targeted sequence. In HCT116 cells more control, non-targeted *Renilla* reporter mRNA was found in fraction 4, but when the reporter mRNA contained a Let-7 target this increase was not seen (compare Fig. 4A and 4D) and there was more reporter mRNA in fraction 1 (“free” mRNA or mRNA complexed with lower molecular weight proteins); these changes were accompanied by a slight increase of luciferase protein level and translation efficiency (Fig.1 and Table 2). In Me45 cells the effect of a Let-7 target was different: the amount of “free” reporter mRNA decreased and it was more abundant in higher molecular weight fractions correlated with an increase in translation efficiency.

**Table 2.**
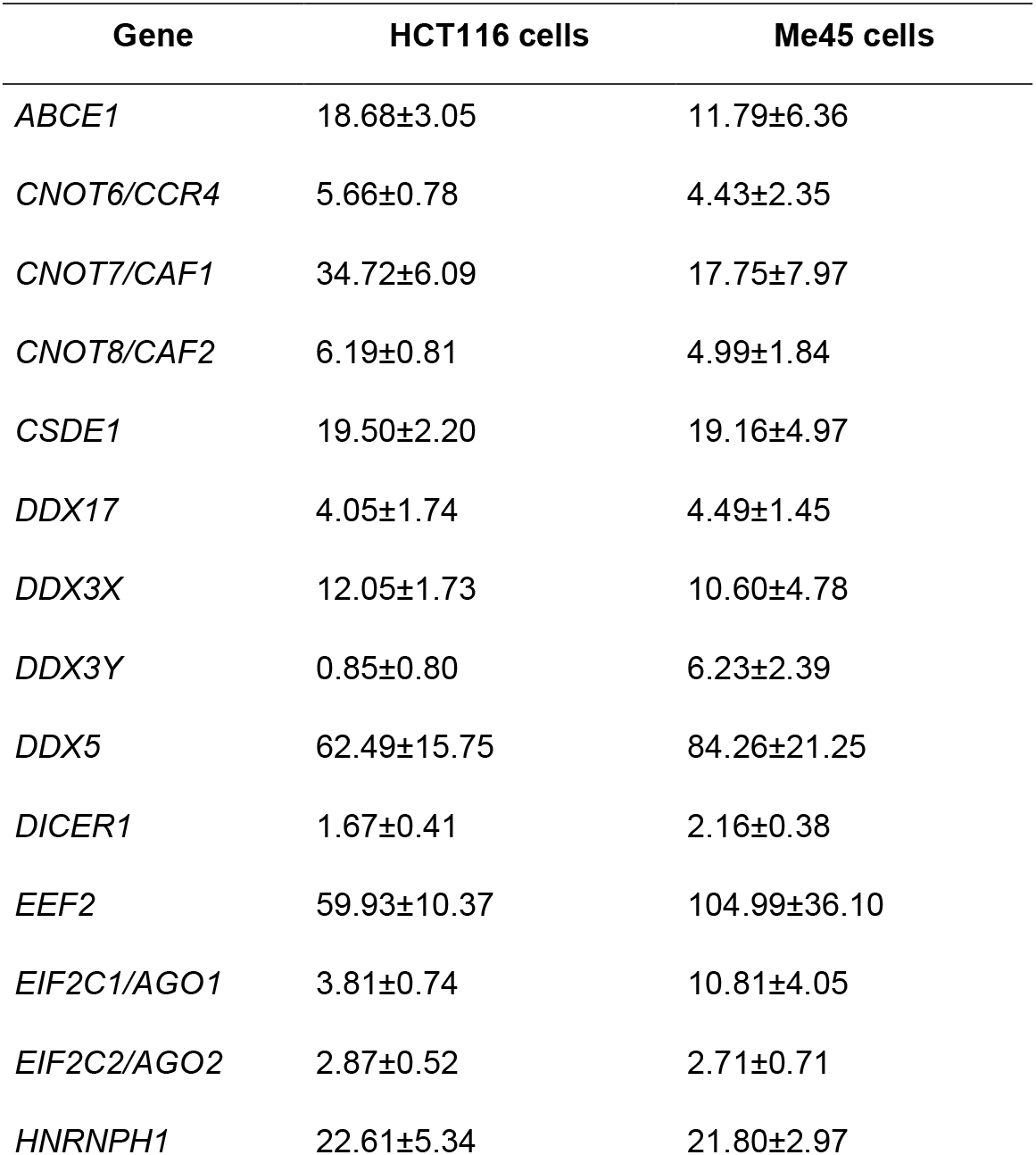
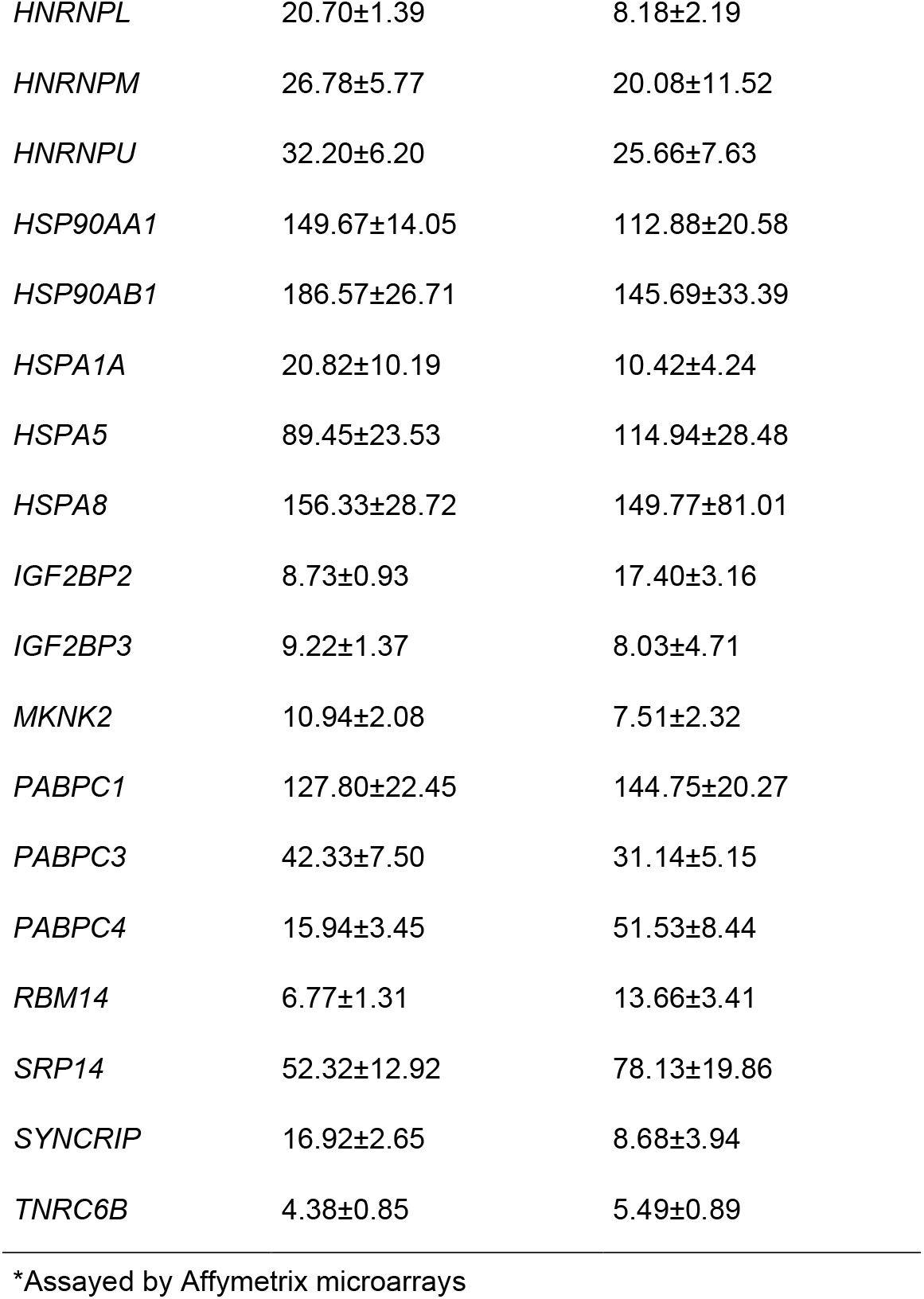
Expression of genes coding for proteins which may be engaged in modulation of translation (mRNA levels*)

| **Gene** | **HCT116 cells** | **Me45 cells** |
| --- | --- | --- |
| *ABCE1* | 18.68±3.05 | 11.79±6.36 |
| *CNOT6/CCR4* | 5.66±0.78 | 4.43±2.35 |
| *CNOT7/CAF1* | 34.72±6.09 | 17.75±7.97 |
| *CNOT8/CAF2* | 6.19±0.81 | 4.99±1.84 |
| *CSDE1* | 19.50±2.20 | 19.16±4.97 |
| *DDX17* | 4.05±1.74 | 4.49±1.45 |
| *DDX3X* | 12.05±1.73 | 10.60±4.78 |
| *DDX3Y* | 0.85±0.80 | 6.23±2.39 |
| *DDX5* | 62.49±15.75 | 84.26±21.25 |
| *DICER1* | 1.67±0.41 | 2.16±0.38 |
| *EEF2* | 59.93±10.37 | 104.99±36.10 |
| *EIF2C1/AGO1* | 3.81±0.74 | 10.81±4.05 |
| *EIF2C2/AGO2* | 2.87±0.52 | 2.71±0.71 |
| *HNRNPH1* | 22.61±5.34 | 21.80±2.97 |
| *HNRNPL* | 20.70±1.39 | 8.18±2.19 |
| *HNRNPM* | 26.78±5.77 | 20.08±11.52 |
| *HNRNPU* | 32.20±6.20 | 25.66±7.63 |
| *HSP90AA1* | 149.67±14.05 | 112.88±20.58 |
| *HSP90AB1* | 186.57±26.71 | 145.69±33.39 |
| *HSPA1A* | 20.82±10.19 | 10.42±4.24 |
| *HSPA5* | 89.45±23.53 | 114.94±28.48 |
| *HSPA8* | 156.33±28.72 | 149.77±81.01 |
| *IGF2BP2* | 8.73±0.93 | 17.40±3.16 |
| *IGF2BP3* | 9.22±1.37 | 8.03±4.71 |
| *MKNK2* | 10.94±2.08 | 7.51±2.32 |
| *PABPC1* | 127.80±22.45 | 144.75±20.27 |
| *PABPC3* | 42.33±7.50 | 31.14±5.15 |
| *PABPC4* | 15.94±3.45 | 51.53±8.44 |
| *RBM14* | 6.77±1.31 | 13.66±3.41 |
| *SRP14* | 52.32±12.92 | 78.13±19.86 |
| *SYNCRIP* | 16.92±2.65 | 8.68±3.94 |
| *TNRC6B* | 4.38±0.85 | 5.49±0.89 |

\*Assayed by Affymetrix microarrays

**Figure 4.**
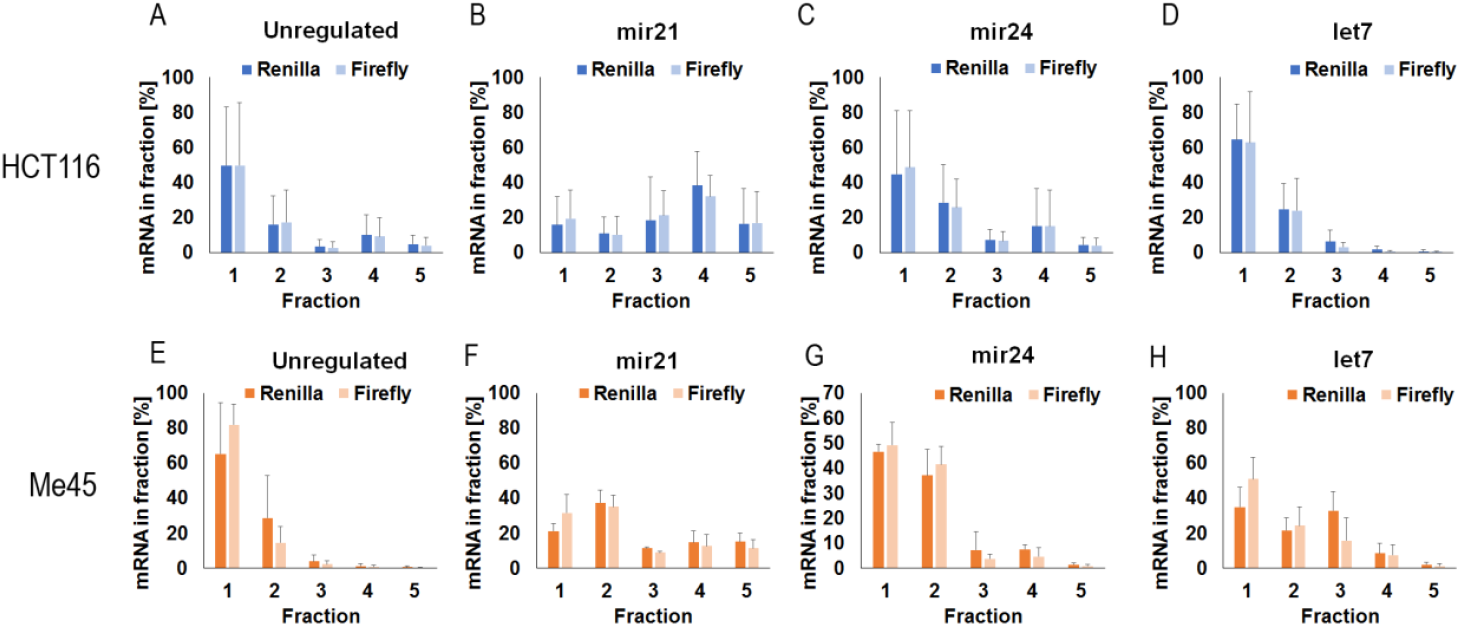
Distribution in sucrose gradients of control and miR21, miR-24, or Let-7-targeted *Renilla* luciferase mRNA and of co-transfected, non-targeted *Firefly* luciferase mRNA from HCT1116 or Me45 cells (upper and lower panels respectively).

The presence of a miR-21 target in mRNA was correlated with less mRNA in fraction 1 and significantly more in heavy fractions in both cell lines; however, some differences between the cell lines were also obvious. In HCT116 cells most of the miR21-targeted mRNA sedimented in the heavier fraction 4, whereas in Me45 cells the peak was in fraction 2 containing complexes lighter than ribosomes. In the case of a miR-24-targeted sequence the distribution of luciferase mRNA in polysomal fractions resembled that of control mRNA in both cell lines, except that in Me45 cells more *Renilla* luciferase mRNA was present in fractions containing higher molecular weight complexes and less in fraction 1.

*Firefly* luciferase mRNA from each cell type also sedimented differently, and its distribution in the gradient changed according to the regulation of co-transfected *Renilla* transcripts. Complexes containing *Firefly* mRNA from both cell types sedimented similarly to those formed by miRNA-targeted mRNAs, in spite of the fact that each miRNA had a specific effect on the distribution of targeted *Renilla* mRNA. The above changes of the properties of complexes containing *Firefly* luciferase mRNA were unexpected, and comparing the changes in *Firefly* mRNA and protein levels with *Renilla* luciferase expression one could conclude that *Firefly* luciferase expression was regulated by some mechanism depending on *Renilla* regulation, even in the absence of miRNA-targeted sequences.

### Influence of anti-miR21, anti-miR24, and different anti-Let7 oligonucleotides on expression of reporter luciferases

To confirm the impact of particular miRNAs on reporter gene expression, we introduced an anti-miRNA oligoribonucleotide for miRNAs which target the *Renilla* gene together with the reporter gene. It is generally assumed that such anti-miR oligonucleotides inhibit miRNA action by forming hybrids with them and thereby prevent their interaction with mRNAs. The effect of anti-miR oligonucleotides for miRNA-21 and miRNA-24 on *Renilla* and *Firefly* luciferase expression are shown in Fig. 5 where expression changes are represented as the decimal logarithm of the ratio of mRNA or protein levels in the presence or absence of an anti-miR-oligonucleotide. As one could expect, addition of an anti-miR-21 to the system with a miR-21 targeted *Renilla* luciferase caused an increase of *Renilla* protein level in both cell types, while the untargeted *Firefly* protein did not change significantly (Fig.5B and D). However, the response of mRNA levels was different in each cell type; in Me45 cells mRNA levels decreased, suggesting that some mRNA molecules are protected from degradation in these cells in the presence of active miR-21.

**Figure 5.**
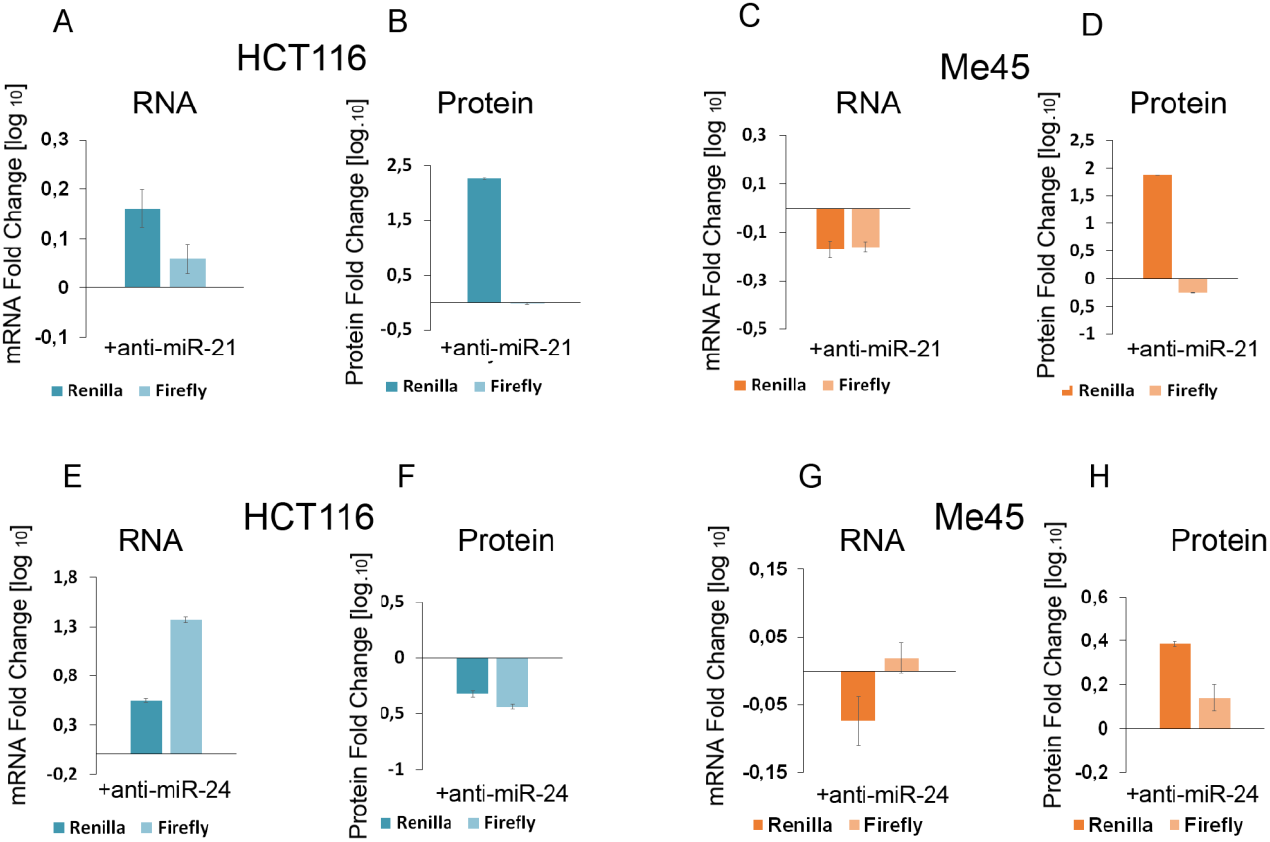
Influence of anti-miR oligonucleotides to miRNA-21 (A,B,C,D), or miRNA-24 (E,F,G,H) on the expression of a miRNA-targeted *Renilla* and co-transfected non-targeted *Firefly* reporter genes. Bar graphs represent the decimal logarithm of the ratio of mRNA (A,C,E,G) or protein (B,D,F,H) levels in the presence or absence of an anti-miR oligonucleotide. Dark blue and dark orange columns show changes of *Renilla* luciferase expression, whereas light blue and light orange show values for *Firefly* luciferase in HCT116 and Me45 cells, respectively.

Anti-miRNA-24 oligonucleotides (Fig.5E-H) appeared to have an opposite effect on protein and mRNA levels in HCT116 and Me45 cells. Their presence also had a visible effect on the levels of the non-targeted *Firefly* luciferase mRNA and protein.

The Let-7 targeted sequence in mRNA is particular in that it may interact with at least 12 different miRNAs belonging to the Let-7 group and encoded by different genes. HCT116 and Me45 cells differ in the levels of these different Let-7 miRNAs (Fig.6C and F). We used anti-miR oligonucleotides to Let-7 family members to see which are most effective in regulation of reporter gene expression (Fig.6). The presence of an anti-miR oligonucleotide to different Let-7 family members had different effects in HCT116 and Me45 cells. In HCT116 cells they caused a decrease of mRNA and protein levels, suggesting that in these cells interactions with any Let-7 has a protective effect on mRNA and that most of such interactions may stimulate translation. In these cells also *Firefly* luciferase expression shows a similar response to different anti-Let-7 oligonucleotides (Fig.6A,B).

**Figure 6.**
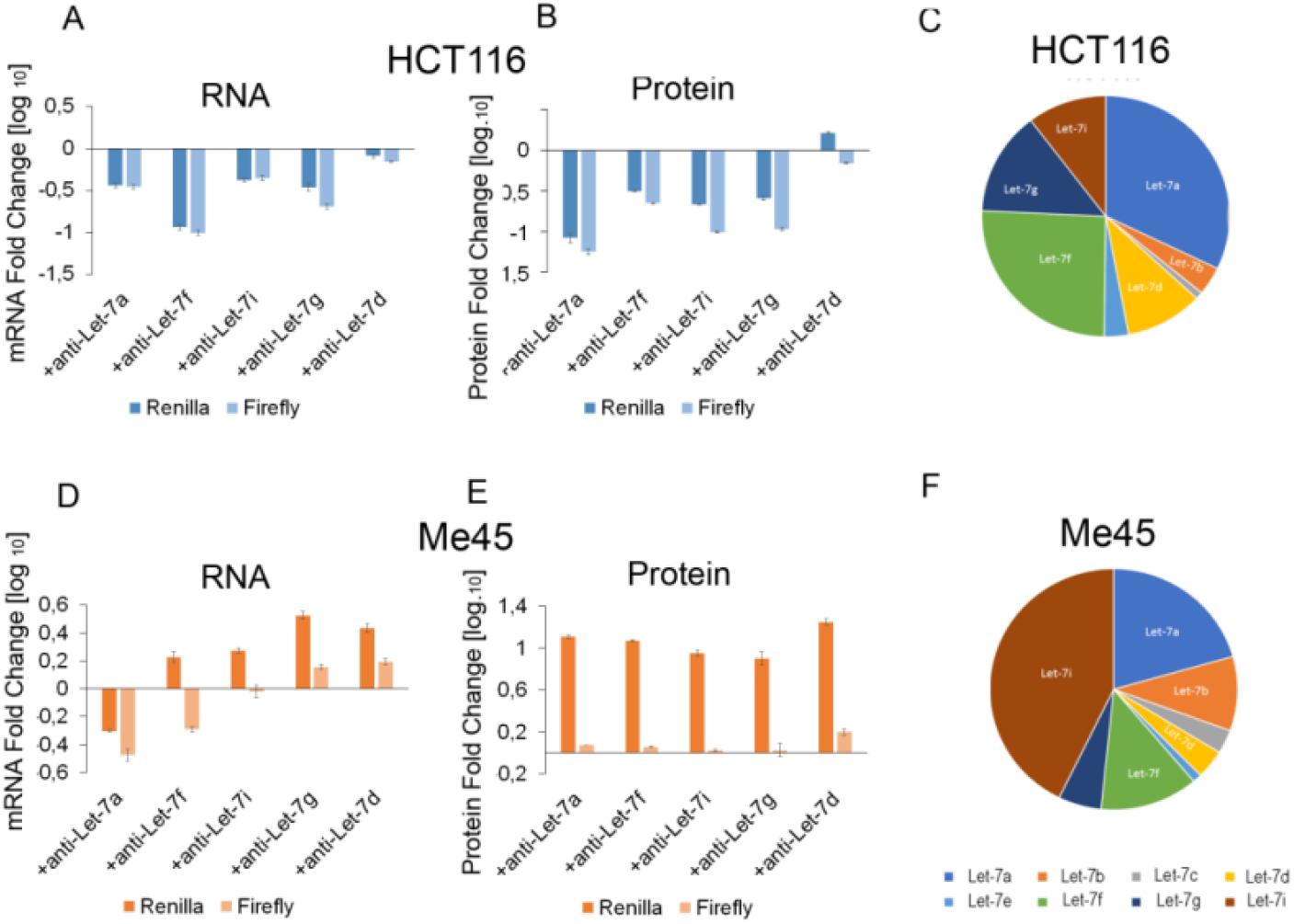
Influence of the inhibition of different Let-7 family members by anti-Let 7miRNA oligonucleotides on *Renilla* and *Firefly* luciferase transcript (A,D) and protein (B,E) levels in HCT116 (A,B) and Me45 (D,E) cells. The relative abundances of the different Let-7 family members in HCT116 and Me45 cells are presented as pie charts (C and F, respectively).

In Me45 cells only the anti-Let-7a oligonucleotides had a similar inhibiting effect on *Renilla* mRNA, the other anti-Let-7s had an opposite effect on mRNA to that seen in HCT116 cells and also the influence on protein level was different. All Let-7 anti-miRs increased the level of *Renilla* luciferase in Me45 cells and (except for anti-Let-7d) decreased it in HCT116 cells. This result suggests differences in mechanisms regulating translation and mRNA stability between cell types and also different participation of Let-7 family members in regulation.

Anti-miR oligonucleotides had similar effects on transcripts of non-targeted mRNAs and their translation in HCT116 cells and caused a decrease in protein level, suggesting that interactions of *Renilla* transcripts with miRNAs somehow protect mRNA and also facilitate translation of non-targeted mRNA.

### Proteins potentially engaged in regulation of translation are differently expressed in Me45 and HCT116 cells

Sucrose gradient centrifugation revealed that a particular mRNA targeted by the same miRNA was localized in different mRNA-protein complexes in Me45 and HCT116 cells (Fig 4). To explore if this could be related to differences in the expression of proteins participating in translation and its regulation, we compared the levels of transcripts for translation initiation factors and other potential regulatory proteins which may interact with AGO or initiation complexes, using Affymetrix microarrays and PCR methods (Fig.7 and Table 3)

**Table 3.**
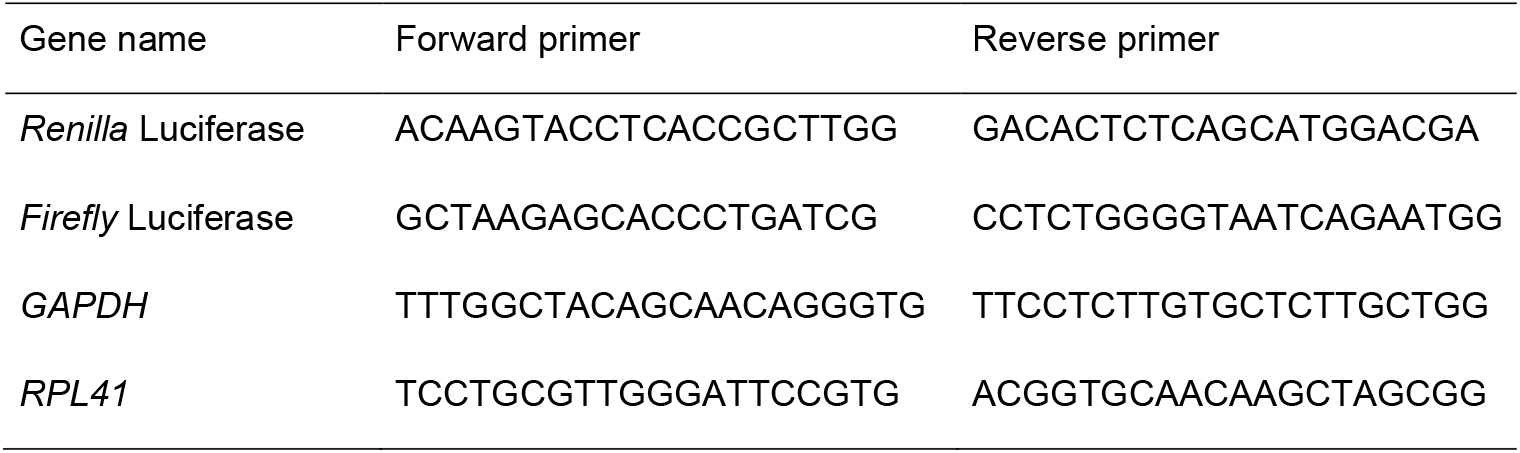
Real Time PCR primers

mRNAs for most initiation factors have similar levels in both cells types and, with some exceptions, the differences do not exceed two-fold. From those which have a higher expression in Me45 cells, EIF1, EIF2S2, some of EIF3, EIF4A1, EIF4EBP1 and EIF6 show the largest difference. In HCT116 cells a strikingly high expression is seen for EIF5A (Fig.7). It is difficult to hypothesize which of these differences could be directly responsible for the differences observed in our experiments, as most translation initiation factors are multifunctional RNA-binding proteins that participate not only in translation regulation but sometimes also in regulation of other cellular processes

**Figure 7.**
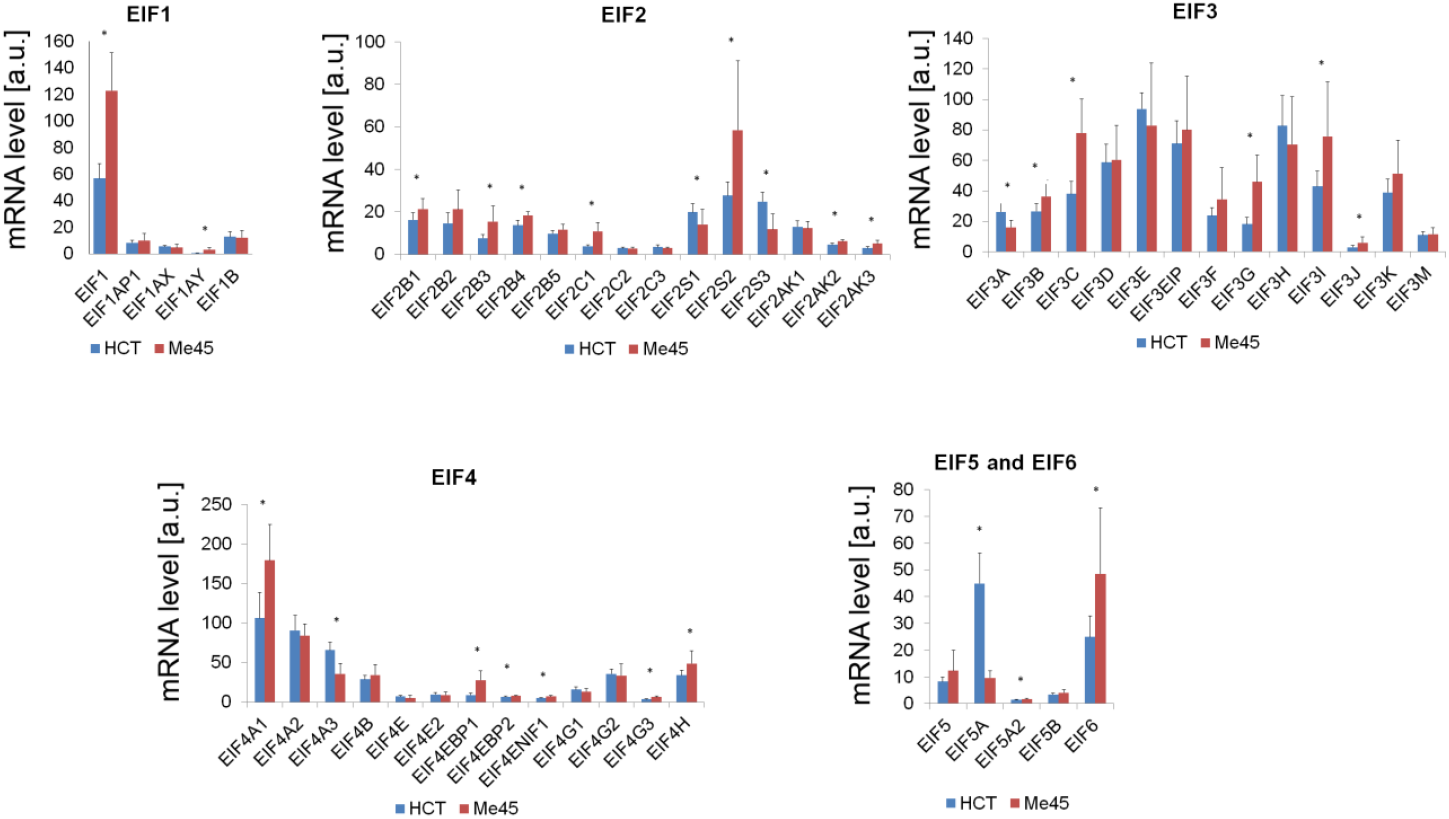
Levels of mRNAs for translation initiation factors in HCT116 and Me45 cells assayed by microarrays (results available in ArrayExpress) normalized to cell numbers used for isolation of RNA. Each panel shows differences in a specific EIF group. The results are presented as mean values ±S.D. of 3 microarray experiments normalized to the number of cells, and * denotes the statistical significance of the difference between HCT116 and Me45 cells (p<0.05)

The efficiency and regulation of translation also depends on the availability and activity of further proteins, which may constitute another layer regulating expression or activity of proteins directly participating in translation. In Table 2 we show examples of the mRNA levels for some of these proteins which were found in complexes regulating translation in both cell types.

Among these proteins, CNOT6, 7, and 8 are subunits of the CCR4-NOT core transcriptional and translational regulation complexes, important for the stabilization, cytoplasmic transport, and deadenylation of polyadenylated mRNAs. CNOT6 (CCR4a) and CNOT7 (CAF1) have RNase activity, and bind to CNOT1 which is a platform able to bind further regulatory proteins (Mathys et al. 2014). ABCE1 is a highly conserved protein required for translation initiation and ribosome biogenesis and is a ribonuclease L inhibitor; it also plays a role in translation termination and ribosome recycling by dissociating ribosomes into large and small subunits (Khoshnevis et al. 2010). Another group of proteins present in Me45 and HCT116 cells that may be crucial for translation regulation are members of the large and highly abundant family of RNA-dependent DEAD-box helicases with ATPase activity (DDXs). Some members of this family participate in liquid–liquid phase separation and are present in RNA-containing phase-separated organelles in prokaryotes and eukaryotes (Hondele et al. 2019). Both HCT116 and Me45 cells show high levels of DDX5 helicase expression. Again, as for initiation factors, one cannot hypothesize which of the differently expressed proteins may be the most important for intercellular differences in miRNA regulation of translation, but taken together these data suggest that differences in the levels of proteins participating in regulating translation initiation could be responsible for the differences in reporter gene expression between HCT116 and Me45 cells.

## DISCUSSION

### Specific miRNA effects in cells of the same type and differences between the effects of the same miRNA in different cell types

In this study, we compared the effects of different miRNAs and posed questions concerning the mechanisms underlying their specificity in the same cell type and the differences between cell types. Reporter genes which differ only in their miRNA-targeted sequences, transfected into the same cell type containing the same sets of proteins which control translation, showed very different expression (Fig. 1); the presence of a miRNA-21 target reduced translation strongly, whereas the presence of a miRNA-24 or a Let-7 target did not change or increased synthesis of the reporter protein in a cell type-specific manner. In different cell types, the same miRNAs induce mRNA-protein complexes that have different sedimentation properties, differ in their influence on mRNA stability and translation efficiency, and react differently to the presence of anti-miR oligonucleotides (Figs. 1, 4, 5 and 6).

These different effects could have many causes. The first may be a difference in the levels of the miRNAs themselves within the cell; in HCT116 cells the miRNA-21 level was about 4 times higher than that of all Let-7s together, and 15 times higher than the level of miRNA-24. In Me45 cells the differences between the levels of miR-21, miR-24 and total (summed) Let-7s were less pronounced (miR21/miR24 ∼ 4 and miR-21/Let-7= 0.8). The higher level of miRNA-21 than of miRNA-24 in both cell types could explain the weaker inhibition of translation by miRNA-24 than miRNA-21, but not the stimulation of expression by miRNA-24 (Fig.1) which needs a different mechanism. The specificity of miRNA binding to various AGO proteins could result in the formation of different complexes and cause differences in the final effect of the miRNA, but published studies rather exclude a specificity of different human AGOs (Yoda et al. 2010; Dueck et al. 2012).

Important factors influencing the operation of the RISC complex are interactions between miRNA and their targets in mRNA, such as the presence and position of mismatches which influence AGO’s action in translation repression and in endo or exo-nucleolytic mRNA degradation (Salomon et al. 2015; McGeary et al. 2019). Our reporter genes differ in the presence and distribution of mismatches; the miR-21 is perfectly complementary, whereas the miR-24 binding site is unpaired on nucleotide 13 and the 3’-most hexanucleotide, and the Let-7 binding site is unpaired on nucleotides 9-12 with every Let-7 family member, and these differences may play a role in eliciting a specific effect. However not all miRNA-specific effects observed here can be directly explained by mismatches and their influence on RISC efficiency, because the effects of the same targets in different cell types are different (see Fig.1, where the influence of targets for miRNA-24 and Let-7 on protein level is opposite in HCT116 and Me45 cells). The intercellular differences in effects of the same miRNAs are most probably caused by differences in formation of RISC-complexes with other proteins resulting from intercellular differences in protein concentrations. In human cells, complexes containing AGO proteins may contain a plethora of other proteins including TNRC6, PABC, HNRNP, HSP, IGFBP family proteins, helicases MOV10 and DEAD box-containing, cold shock domain-containing proteins, RBM14, TP53, translation elongation factors, and ribosomal proteins as shown by immunoprecipitation (Dueck et al. 2012; Landthaler et al. 2008; Kakumani et al. 2020; Meister et al. 2005). In *C. elegans*, the same miRNA is found in RISC complexes of different composition in different cell types (Dallaire et al. 2018). RISC complexes with different compositions may co-exist in the same cell, the frequencies of those which contain or not helicases, nucleases or nuclease inhibitors depending on the availability of these components at any given time. Further, proteins in AGO and TNRC6 complexes may be modified post-translationally by prolyl-hydroxylation, phosphorylation, ubiquitination, or poly-ADP-ribosylation, which may alter their interactions and influence miRNA activity at global or specific levels (Munakata et al. 2021). Different protein-RISC complexes may degrade mRNA or inhibit translation with or without mRNA degradation, or sometimes even stimulate translation depending on their composition (for example through interactions of PABP-EIF4F). In our experiments, the presence of sequences targeted by Let-7 did not lead to mRNA degradation in HCT116 cells and was even associated with protection, because the addition of oligonucleotides complementary to various representatives of the Let-7 group caused a decrease in the level of the reporter gene mRNA (Fig. 6C). RISC associated with TNRC6-PAN or TNRC6-CCR4-NOT promotes mRNA deadenylation and degradation, but the presence of other proteins such as some helicases involved in the formation of P-bodies or stress granules could lead to the formation of nucleation centers and translation-blocking complexes.

The operation of RISC may depend not only on the nucleotide sequence of the target but also on the conformation of the entire mRNA molecule, which always tends to fold and adopt spatial structures stabilized by complementary interactions. The involvement of the seed sequence in the formation of a double-stranded structure within the mRNA chain may make its interaction with RISC difficult. In Fig.8, we compare the conformation of non-complexed “naked” Renilla luciferase transcripts containing targets for miRNA-21 and miRNA-24 obtained using the RNAStructure (Reuter and Mathews 2010) software.

**Figure 8.**
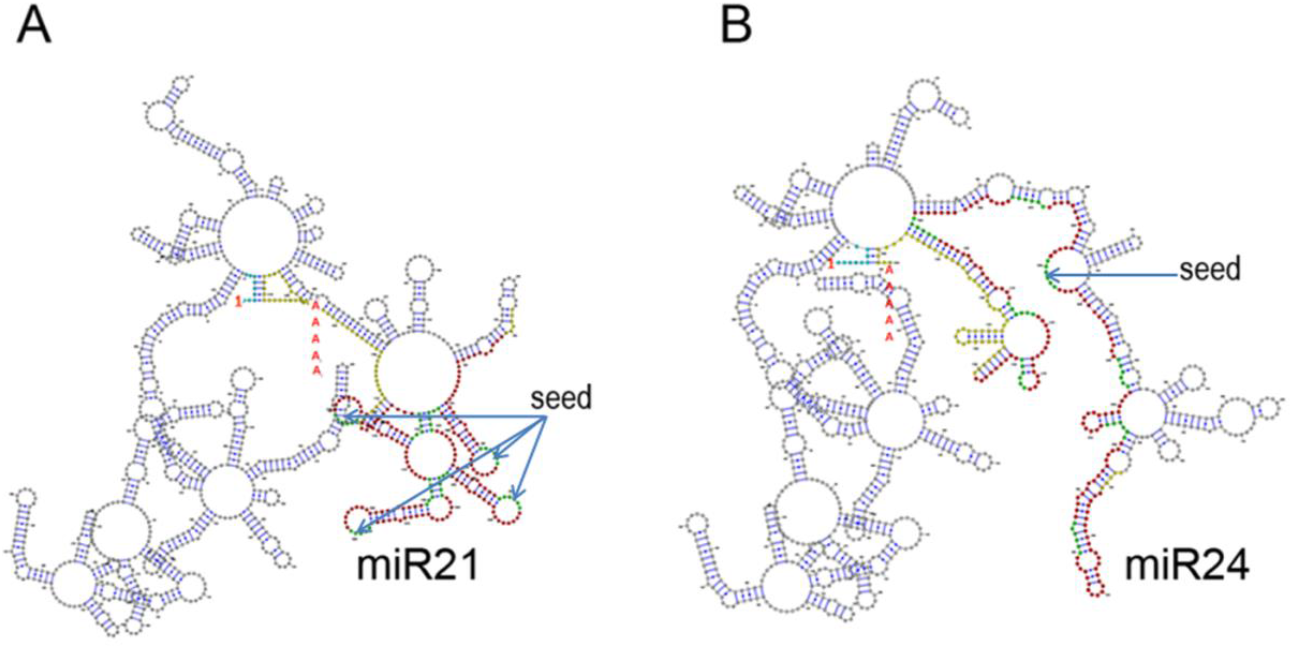
Most probable structures of *Renilla* mRNA containing 8 repeated inserts complementary to miR-21 (A), or miR-24 (B), predicted by the program RNAstructure (Reuter and Mathews 2010) and visualized by the tool VARNA GUI (Darty et al. 2009). Black or colored dots represent nucleotides; the first is marked by a red number 1 and the 5’UTR nucleotides are blue; 3’UTR nucleotides are yellow; miRNA-targeted inserts are red, and seeds which bind RISC are green, the beginning of the poly(A) sequence is marked by a red letter A, the arrows indicate the seeds that do not form complementary bonds.

Conformations with the highest stability (the lowest energy) differ in the involvement of the “seed” sequences in the formation of double-stranded structures. Four seeds from eight miR-21 targets tend to stay in a single strand conformation, whereas in transcripts with miR-24 targets only one seed appears to be single stranded. However, one has to remember that the transcript conformation within the cell may significantly differ because of interaction with cellular proteins. In summary, the differences in function between individual miRNAs in the same cells most probably depend on the nucleotide sequence of the target present in mRNA and the spatial conformation of the whole transcript which may determine the composition of the RNP complex.

The differences between cell types in the effects (action) of the same miRNA are due to differences in the levels of this miRNA, but also to differences in the concentration and distribution of proteins capable of complexing with RISC, which is best seen in experiments with anti-miR oligonucleotides. Limiting the action of miRNA-21 by anti-miR in both cell types goes hand in hand with an increase in the level of protein, which is consistent with the concept of translation inhibition. However, in the case of mRNA, the effects in HCT116 and Me45 cells are opposite; in the former the mRNA level increases but in the latter it decreases in the presence of the same anti-miR (Fig. 5). This suggests that despite similar effects on translation, the miRNA-21 initiated complexes have different composition and interact differently with mRNA in each cell type and that in the case of anti-miR24 and different Let-7 group members the opposite effects in both cell types also concern protein levels (Fig.5 and 6).

### Pitfalls in normalization of results for miRNA-targeted gene expression to those for a non-targeted gene

In our transient transfection experiments, we observed that the presence of a target for miRNA on *Renilla* luciferase transcripts affected the expression efficiency of a non-targeted *Firefly* luciferase genes. Conventionally, in studies with two co-transfected plasmids the observed expression of a miRNA-targeted gene is “normalized” to the expression of a second gene regulated only by a promoter, in order to correct for differences of transfection efficiency. Here we used a different strategy to analyze the results of studies of the influence of miRNAs on the expression of targeted reporter genes cloned in psiCHECK-2 plasmids; we analyzed separately the expression of both the co-transfected miRNA-regulated and the non-regulated gene, calculating and comparing the levels of mRNA and protein per single cell. This strategy revealed some unexpected features which are masked in the conventional normalization method; insertion of a miRNA target into the *Renilla* luciferase gene caused changes not only of its own transcript and protein levels, but also of those of a co-transfected but non-targeted *Firefly* reporter gene (Fig. 1). Moreover the targeted and non-targeted mRNAs are in similar protein-mRNA complexes as seen by sucrose gradient centrifugation, but which differ depending on the type of miRNA target on the *Renilla* gene (Fig.4), and anti-miRNA oligonucleotides which theoretically should not influence the expression of mRNAs which are not regulated by these miRNAs affect also the expression of the *Firefly* gene (Figs.5 and 6). Altogether, such results suggest that targeted and non-targeted mRNA molecules participate in the same regulation process initiated by a miRNA. We hypothesize that such common regulation could be based on a process similar to stress granule or P-body formation by creation of nucleation centers and further liquid-liquid phase transition, as proposed for the formation of P-bodies or stress granules by a combination of RNA-RNA, protein-RNA and protein-protein interactions (Protter and Parker 2016; Banani et al. 2016; Kedersha et al. 2016; Panas et al. 2016); their formation depends on trans RNA-RNA interactions which may be induced by activation of DEAD box helicase and its ATPase activity (van Treeck et al. 2018; Mugler et al. 2016; Khong et al. 2017).

Connections between P-bodies and AGO-induced inhibition of translation and mRNA degradation are known since many years, and miRNA-directed inhibition of translation appears to be necessary for P-body or stress granule formation (Jakymiw et al. 2005; Eulalio et al. 2007). It was also found that trans RNA-RNA interactions induce the core of nucleation centers leading to liquid-liquid phase transition (Garcia-Jove Navarro et al. 2019). In our experiments inhibition of translation is accompanied by formation of polysome-size complexes (Fig. 4) which are significantly smaller than bodies visible by fluorescence microscopy. Nevertheless, the presence of miRNA-targeted and non-targeted mRNAs in these complexes would suggest the existence of nucleation centers whose further growth is somehow inhibited (for example by the presence of NOT1 protein which inhibits formation of P-bodies (Sachdev et al. 2019) and could be attracted by RISC). The decreased expression level of the untargeted *Firefly* luciferase gene when the transcript of a co-transfected *Renilla* luciferase gene contains a miRNA target, could be explained by such a mechanism of compact complex formation.

### Can 5’-3’ trans interactions between miR-targeted and non-targeted mRNAs explain stimulation of translation?

To explain the increased expression of non-targeted mRNAs in the presence of targeted ones (Fig.1D) we propose a further type of interaction which potentially could be influenced by RISC. The importance of mRNA circularization for efficient translation has been widely discussed (for example (Jackson et al. 2010; Vicens et al. 2018; Wells et al. 1998; Alekhina et al. 2020)), and is believed to be responsible for the more frequent re-initiation on circularized compared to linear mRNA (Wells et al. 1998; Gallie 1991; Alekhina et al. 2020). The proximity of the 5’ and 3’ ends of the same RNA molecule suggested by the structure of naked mRNA ((Vicens et al. 2018) and Fig.8)), if persisting inside the cell, would favor their interaction, and Filipowicz and his group (Filipowicz et al. 2008; Pillai et al. 2007; Mathonnet et al. 2007) have proposed that circularization of mRNA molecules can occur through interaction of an EIF4F complex on the 5’ end with PABP proteins bound to the 3’ end, but is prevented by binding of RISC. On the other hand, in resting cells there are also examples of stimulation of the translation of some cellular and reporter genes by the interactions of RISC containing AGO2 with miRNA and the FXR1 protein, which was explained by the formation of a complex joining the 5 ‘and 3’ ends of the mRNA which would facilitate the re-initiation of translation (Vasudevan et al. 2008; Truesdell et al. 2012; Vasudevan et al. 2007). Such RNA circularization by protein complexes is usually assumed to occur only in *cis*, but *trans* 5′–3′ interactions between different mRNA molecules are not experimentally excluded. Following an earlier idea (Filipowicz et al. 2008), we propose that binding of RISC to the 3’ end of an mRNA is likely to create a hindrance to circularization of single transcript but in some types of complexes may facilitate *trans* 3’–5’ interactions of mRNA molecules, which could confer similar advantages for activation of translation initiation as in circularized single molecules. This model of inter-molecular activation of translation would be consistent with the results of Archer and coworkers (Archer et al. 2015) who probed a closed-loop model of mRNA translation and found no correlation between the amounts of EIF4F components and of PABP bound to the 5’ and 3’ ends of the same mRNA molecule; these results for 16 genes could be explained by trans 3’-5’ interactions with other types of mRNAs and suggest that the *trans* PABP-EIF4F interactions which we propose may be quite common.

Our results are consistent with a model where translation of a population of specific mRNA molecules is regulated by the formation of several alternative and different types of mRNA-RISC complex which contain single closed circular as well as multiple mRNA molecules, so that the levels of different RNA and protein products measured experimentally is the net result of these processes.

The suggested close vicinity of 5’ and 3’ ends of an mRNA resulting from intrinsic RNA properties (Vicens et al. 2018) would favor the preponderance of structures containing single mRNA molecules, but interaction of mRNA 3’ ends with some miRNPs could change this situation by creating opportunities for further interactions leading to mRNA degradation or nucleation centers or trans 3’ - 5’ complex formation between of poly(A) - bound PABP and EIF4F located on another mRNA molecule. At present, these models are mainly conceptual, and more detailed experimental studies of the relevant RNA-RNA and protein-RNA interactions are needed. New methods which allow to explore the multiple conformations and interactions of RNAs in living cells (Ziv et al. 2018) create new possible directions for further investigations.

## MATERIALS AND METHODS

### Cell lines

HCT 116 cells (human colorectal cancer, ATCC CCL-247) and Me45 cells (melanoma, Oncology Center, Gliwice described in (Kramer-Marek et al. 2006)) were grown in standard conditions in DMEM/F12, (PAN Biotech, Aidenbach, Germany) supplemented with 10% fetal bovine serum (Eurx, Gdansk, Poland) and penicillin-streptomycin (Sigma-Aldrich, St. Louis, USA), at 37ºC in a humidified atmosphere with 5% CO_2_.

### Plasmids and transfection

Cells were grown to 70-80% confluency before transfection (about two or three days). Cells were transfected with psiCHECK-2 plasmids (Promega, Madison, USA) containing two luciferase genes, a reference *Firefly* gene and a reporter *Renilla* gene containing eight tandem repeats of target sequences for miRNAs Let-7, miR-21 and miR-24 in their 3’UTRs. The Let-7 target sequence contained the motif TCGAGACTATACAAGGATCTACCTCAG with average complementarity of 71.75% to various mature let-7 family members, and the miR-21 target TCAACATCAGTCTGATAAGCTAAA, was 100% complementary to the mature miR-21 sequence; the two last AAs form a spacer to limit complementarity for nonspecific binding. The miR-24 target sequence contained the motif ATACGACTGGTGAACTGAGCCG, 68% complementarity to the mature miR-24. psiCHECK-2 with Let-7 sequence targets was a kind gift from Martin Simard, Laval University, Quebec. Sequence synthesis, insertion, and verification were performed by BLIRT (Gdansk, Poland). The unmodified plasmid was used as an unregulated control. Transfection was performed with branched polyethylenimine (Sigma-Aldrich, St. Louis, USA) according to the supplier’s protocol using 1 μg of plasmid per well in 12-well plates or 8 μg for a T75 flask and cells were harvested 24 h after transfection.

### nti-microRNA oligonucleotides (anti-miRs)

Seven miRNA-inhibiting oligonucleotides (anti-miRs) were purchased from Integrated DNA Technology (Coralville, Iowa, USA). Their sequences were complementary to those in miRbase (hsa-miR-21-5p: MIMAT0000076; hsa-miR-24-3p: MIMAT0000080; hsa-let-7a-5p: MIMAT0000062; hsa-let-7d-5p: MIMAT0000065; hsa-let-7f-5p: MIMAT0000067; hsa-let-7g-5p: MIMAT0000414; hsa-let-7i-5p: MIMAT0000415). Twenty pmol per well in 12-well plates were co-transfected with the appropriate plasmid.

### Extraction and assay of RNA

Total and polysomal mRNA were assayed by RT-qPCR. RNA was extracted with a Total RNA Mini kit (A&A Biotechnology, Gdynia, Poland), and reverse transcription was done with a NG dART kit (Eurx, Gdansk, Poland) using both oligo(dT) and random hexamers according to the supplier’s protocol. qPCR was performed on a CFX96 Touch Real Time PCR System (BioRad, USA) with the RT PCR Mix SYBR® A kit (A&A Biotechnology, Gdynia, Poland).

The numbers of reporter transcript molecules in cells were calculated with the help of calibration curves constructed for each reporter gene on the basis of DNA PCR performed with known numbers of plasmid DNA molecules containing analyzed gene.

### Luciferase assays

Luciferases were assayed by their activity in oxidizing luciferin with emission of luminescence using the Dual-Luciferase Reporter Assay System (Promega, USA) according to the producer’s protocol. Luminescence was measured with an Infinite F200 Pro microplate reader (Tecan, Männdorf, Switzerland) on all-white flat-bottom 96 well plates (Falcon/ Corning, USA). Activities are expressed as arbitrary units (a.u.)

### Sucrose gradient centrifugation

Three hours before harvesting cells the culture medium was changed for fresh medium. Cells were incubated for 5 min with cycloheximide (Sigma-Aldrich, St. Louis, Missouri, USA cat. num.:C1988-1G) (100 ng/ml), then washed with ice-cold PBS (PAN Biotech cat. num.: P04-36500, Aidenbach, Germany) containing cycloheximide (100 ng/ml), detached by scraping, and immediately lysed or frozen in liquid nitrogen and stored for up to 6 weeks before lysis. Lysis buffer was polysome buffer (10 mM KCl, 2 mM MgCl_2_, 20 mM Tris-HCl pH 7.6, 1 mM DTT, 100 ng/ml cycloheximide) supplemented with 0.5% Triton X-100 and 2.5% glycerol, used ice cold. Extracts were centrifuged at 16 000 x g for 10 min and the supernatant was layered on a pre-formed 15% - 45% sucrose gradient in polysome buffer in a 12 ml polyallomer tube. The gradient was centrifuged in a SW41ti rotor in an Optima XPN-100 Ultracentrifuge (Beckman Coulter, USA) at 4**º**C for 4 h and 100 two-drop fractions were collected into Eppendorf tubes by piercing the tube bottom. The RNA content of fractions was measured on a NanoDrop 2000 (ThermoScientific, USA) at 260 nm, and on the basis of the absorbance profile succeeding fractions were combined to create 5 larger fractions which should contain (from the top) free RNA, small ribosome subunits, monosomes with large ribosome subunits, light polysomes, and heavy polysomes and these fractions served for further analyses.

### Calculation of cell numbers serving for assays

The number of cells used in experiments were calculated from the total RNA in samples and the RNA content per cell, by measuring total RNA from 100 000 cells spectrophotometrically and counting cells by microscopy. The RNA/cell from triplicate assays was 20.97 ± 5.66 pg for HCT116 cells and 35.27 ± 5.78 pg for Me45 cells.

### Microarray analyses

To examine the influence of the levels of different eukaryotic initiation factors (EIFs) and proteins engaged in regulating translation, we compared the levels of their transcripts using culture conditions and Affymetrix microarrays described in our earlier studies (Herok et al. 2010; Rzeszowska-Wolny et al. 2009). The results are available in the ArrayExpress database under accession number E-MEXP-2623 (Athar et al. 2019). miRNA levels were estimated by Agilent microarrays (G4870A SurePrint G3 Human v16 miRNA 8×60k) and are available in the ArrayExpress database under accession number E-MTAB-5197 (Athar et al. 2019). All data are MIAME compliant. Microarray data quality was assessed using the simpleaffy Bioconductor package (Wilson and Miller 2005). Raw HG-U133A microarray data from two experiments on both cell lines were processed using Brainarray EntrezGene specific custom CDF (v22) (Dai 2005) in R using the RMA algorithm implemented in the affy Bioconductor library (Gautier et al. 2004). Differentially expressed genes were identified using limma with a q-value correction for multiple testing (Camps et al. 2014; Ritchie et al. 2015). Genes coding for proteins engaged in regulating translation were identified using GO terms (Yoda et al. 2010; The Gene Ontology Consortium 2019). To compare the levels of transcripts in HCT116 and Me45 cells, microarray results were normalized to the number of cells used for the assays.

### RNA double stranded conformation

The conformation of mRNAs for *Renilla* luciferases was calculated by the RNAStructure program (Reuter and Mathews 2010) and visualization was by the VARNA GUI tool (Darty et al. 2009).

### Statistical tests

Results are presented as mean +/-SD from at least three separate assays performed on cells from the same passage. All calculations were performed using Microsoft Office 2016. Significances were determined by T-tests.

## ACKNOWLEDGMENTS

The authors cordially thank Ronald Hancock for discussions and revision of the manuscript. Krzysztof Biernacki is thanked for help in establishing the methods and performing the first transfections.

